# A Multi-Granularity Approach to Similarity Search in Multiplexed Immunofluorescence Images

**DOI:** 10.1101/2023.11.26.568745

**Authors:** Jennifer Yu, Zhenqin Wu, Aaron T. Mayer, Alexandro Trevino, James Zou

**Affiliations:** Enable Medicine & Department of Computer Science, University of Toronto; Enable Medicine; Department of Biomedical Data Science Stanford University

## Abstract

Due to the rapid increase and importance of multiplexed immunofluorescence (mIF) imaging data in spatial biology, there is a pressing need to develop efficient image-to-image search pipelines for both diagnostic and research purposes. While several image search methods have been introduced for conventional images and digital pathology, mIF images present three main challenges: (1) high dimensionality, (2) domain-specificity, and (3) complex additional molecular information. To address this gap, we introduce the MIISS framework, a **M**ulti-granularity m**I**F **I**mage **S**imilarity **S**earch pipeline that employs self-supervised learning models to extract features from mIF image patches and an entropy-based aggregation method to enable similarity searches at higher, multi-granular levels. We then benchmarked various feature generation approaches to handle high dimensional images and tested them on various foundation models. We conducted evaluations using datasets from different tissues on both patch- and patient-level, which demonstrate the framework’s effectiveness and generalizability. Notably, we found that domain-specific models consistently outperformed other models, further showing their robustness and generalizability across different datasets. The MIISS framework offers an effective solution for navigating the growing landscape of mIF images, providing tangible clinical benefits and opening new avenues for pathology research.

## 1 Introduction

Multiplexed Immunofluorescence (mIF) is a biological imaging technique that allows the detection and localization of many antigens in a single tissue sample (1). By using multiple fluorescence-labeled antibodies, it provides a comprehensive view of cellular interactions and morphology, making it invaluable for medical research and clinical diagnostics (2) (3). Recently, there has been a significant surge in the volume of mIF data within the field of spatial omics. This has made it increasingly important to streamline the process for pathologists to access and search through available mIF images for diagnostic analysis and research. Traditionally, stored Hematoxylin and Eosin (H&E) images are tagged along with metadata and pathologists could search specific keywords to find relevant images (4)(5). However, this text-based search is unable to capture the complex anatomical and morphological features present in the pathology images, thus limiting access to more comprehensive information (6) (7) (8). Additionally, due to the challenges posed by large image sizes, complex tissue structures, and cellular heterogeneity, it becomes time-consuming to label each individual cell image (9)(10). The absence of such labeling further complicates text-based searches across H&E databases. Similar challenges apply to mIF data as well, with additional challenges of data high-dimensionality and molecular features. Given the expanding volume of mIF data, it is no longer feasible for pathologists to manually review every image in the database. Therefore, these challenges underline the need for an efficient image-to-image search pipeline in mIF data.

The recent advances in artificial intelligence, particularly in the domain of self-supervised learning (11)(12)(13)(14), have demonstrated considerable promise in extracting meaningful feature embeddings from medical images, which have proven effective in diverse image-based tasks such as classification (15)(16), and similarity search (17). State-of-the-art (SOTA) models(17; 18; 19) have demonstrated superior performances in H&E image-to-image retrieval tasks. However, there has yet to be a specialized image search pipeline for mIF images. Although mIF image is also microscopic, it is more challenging, complex and domain-specific due to its high dimensionality and additional unique molecular information. Like H&E images, it is necessary to segment mIF images into smaller patches for any image search tasks. However, the goal extends beyond merely identifying similar patches; there is also a need to aggregate the patch-level information to perceive similarities at a higher level such as patients. This necessitates a similarity search method capable of finding similar samples across multiple levels of granularity, which offers tangible clinical benefits. For instance, the ability to identify similar cellular structures at the patch level can greatly speed up the data labeling process for pathologists. Moreover, compiling search results from patch-level data to regions or even entire patient profiles enables a more holistic clinical analysis such as retrieving rare diagnoses or tissue states. The aggregated approach could be crucial in determining patient outcomes such as HPV status or survival status, selecting specific cohorts and developing new biomarkers. Beyond these immediate applications, the capability to search for similar samples opens up new avenues for clinical research providing pathologists with novel methods for investigating specific regions of interest within tissue specimens. To address the demands of various applications, we introduce MIISS for **M**ulti-granularity m**I**F **I**mage **S**imilarity **S**earch, utilizing self-supervised learning models and entropy-based aggregation methods. Through benchmarking various feature generation approaches and testing foundational models, our evaluations show accurate retrieval of tissue morphology and patient outcomes across both patch and patient-level tasks in multiple datasets. We also found that domain-specific models exhibited superior performance and robustness across diverse datasets.

## 2 Related Works

While no research to date explicitly tackles the retrieval task for spatial omics, there have been notable studies focused on image retrieval in H&E data. One such method Yottixel (19) employs a unique approach that begins by constructing a “mosaic” of representative patches from Whole Slide Images (WSIs). These patches are encoded using features extracted by DenseNet (20), which are then converted into barcodes for search functionality. In a similar vein, the SISH (18) method also assembles a mosaic but incorporates a pre-trained vector quantized variational autoencoder (21) to establish an indexing system. Both SISH and Yottixel transform WSIs into mosaics, generate pertinent indices and features, and apply specialized search algorithms to identify similar entries. In parallel, recent efforts on large scale H&E language-image joint pretraining allow image retrieval with both text and image inputs. One representative method, the Pathology Language–Image Pretraining (PLIP) (17), used H&E data collected from medical Twitter to develop a versatile image and language foundation model for pathology. Similar to PLIP, another multimodal method CONCH (22) achieved higher performance through contrastive training using a diverse set of H&E image-caption pairs.

## 3 Method

The MIISS framework is composed of three core components: 1) generate feature embeddings from mIF image patches, 2) retrieve relevant patches, and 3) aggregate at the patient level based on the results of patch retrieval. For a comprehensive overview of the methodology, please refer to Figure 1.

**Figure 1.**
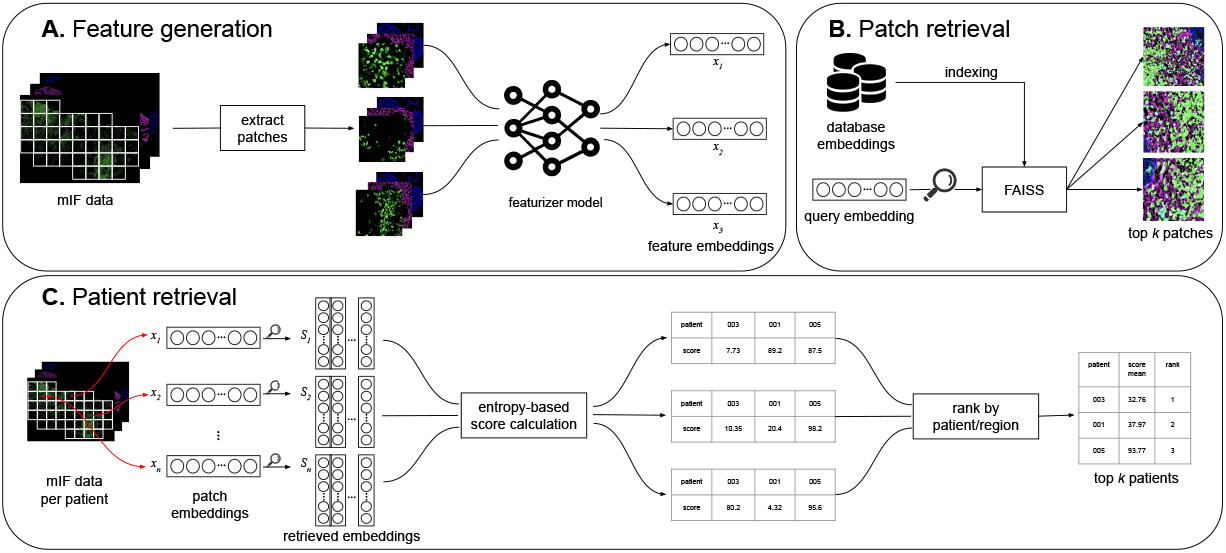
Method Overview: A. feature embeddings generation, B. patch-level retrieval, C. patient-level retrieval

### 3.1 Feature Generation

To generate feature embeddings, we first broke images of full regions (corresponding to tissue micro-array cores) into smaller image patches to capture local molecular localization and morphological features. Subsequently, we applied two specialized transformations: random colorization and feature fusion, to transform each multi-channel patch into one or multiple 3-channel RGB images. Once each multi-channel patch is transformed, we obtain its corresponding feature embeddings.

#### 3.1.1 Patch Generation

Each region contains a tissue micro-array core. After acquiring mIF images of the region, we first identified the upper and lower bounds of fluorescence signals based on image histograms and applied min-max normalization to each channel. Then to acquire image patches containing local features, we employed a sliding window mechanism to draw square bounding boxes with side lengths of 256 pixels (around 100*μ*m) on the region. The stride of bounding boxes was set to 64 pixels to ensure exhaustive coverage. We additionally used a convolutional neural network-based tissue foreground predictor to exclude bounding boxes that have less than 90% tissue coverage. Image patches corresponding to the resulting set of bounding boxes were extracted and saved for feature generation.

#### 3.1.2 Featurizer

We evaluated the performance of three image feature extraction models: DinoV2 (14) a SOTA self-supervised learning model trained on natural images; the PLIP image encoder (17), a visual-language foundation model specifically trained on H&E images; and the ResNet encoder (23), a CNN model trained with natural images from ImageNet. These models were selected as they represent a diverse set of pre-trained models, capturing both general and domain-specific feature extraction capabilities. Additionally, we incorporated a naive averaging method that computes the mean value for each biomarker channel within the patch, serving as a baseline for comparison.

#### 3.1.3 Multi-channel transformations

Given that our selected feature extraction models were originally trained with 3-channel RGB images, a conversion of the multi-channel mIF images to a 3-channel format is necessary. It is important to note that most mIF images typically contain a substantial number of channels, often ranging from 40 to 100 channels. In later sections, *“random colorization”* or *“fusion”* is added after the model name to represent the use of random colorization or feature fusion method.

##### Random Colorization

One approach is a straightforward compression of multi-channel mIF images to 3-channel RGB images by randomly assigning different RGB values to each biomarker. This approach retains the morphological features of tissues presented in different biomarkers and it allows straightforward integration and comparison between regions profiled with different sets of biomarkers. However, due to the fact that it removes identities of the channels, it might cause confusion between biomarkers and introduce extensive noise, potentially compromising the ability to differentiate between patches based on key signature biomarkers.

##### Feature Fusion

An alternative approach partitions the entire set of biomarkers into distinct subsets, with each containing three random biomarkers. Each subset is independently processed through the image encoder, and the feature embeddings generated from subset are concatenated to form a comprehensive feature vector, thereby preserving the multi-dimensionality of the original mIF data. When comparing images profiled with different biomarkers, only overlapping biomarkers will be used. One drawback of the approach is its significant computational demand and extended processing time.

### 3.2 Similarity Search

#### 3.2.1 Patch-level search

After generating feature embeddings, the distance between each query embedding (**x** *∈* ℝ^*n*^) and all database embeddings(**D** *∈* ℝ^*J×n*^) are computed. For a given query sample, its distance to the *j*-th entry in the database is determined using the Euclidean (*l*_2_) distance between the query embedding **x** and the database embedding **d**_*j*_:

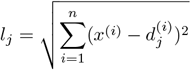

where *x*^(*i*)^ represents the *i*-th component of the embedding vector **x**.

We retrieve the top *m* most similar patches in **D** based on the Euclidean distances demonstrated above. The Euclidean distance was chosen because the magnitude of feature embeddings is significant in this case for finding similar patches. FAISS (Facebook AI Similarity Search, (24)) is employed to calculate all Euclidean distances between embeddings, as it is well-suited for large-scale high-dimensional similarity search due to its efficiency, scalability, and flexibility.

#### 3.2.2 Patient search

To retrieve entries on a patient level, we first extract the embeddings for all patches from the query patient, defined as **X** = [**x**_1_, **x**_2_, …, **x**_*P*_ ], **X** *∈* ℝ^*P ×n*^, where *P* is the number of query patches. For the *p*-th query patch, we define its *m* most similar entries as *𝒮*_*p*_ = [*s*_1_, *s*_2_ …, *s*_*m*_], *𝒮*_*p*_ *∈* ℝ^*m*^.

The corresponding distances between the *p*-th query patch and its *m* most similar database patches are denoted as 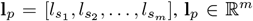 and their respective patient identifiers are 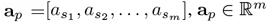.

Among all *K* patients in the database, the mean distance from the *p*-th query patch to the *k*-th patient Pat_*k*_ can be calculated as:

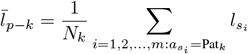

where 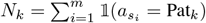 is the number of hits on patient Pat_*k*_ from the *p*-th query patch. We hence can derive the occurrence frequency of Pat_*k*_ as 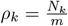, where apparently 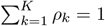

Based on the frequencies, we calculate the entropy of the retrieval results of the *p*-th query patch as:

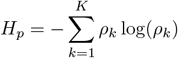

The distance score from the *p*-th query patch to the *k*-th patient in the database is calculated as:

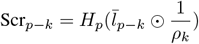

The final score for the distance between the query patient and the *k*-th patient in the database can be calculated as the aggregation of all query patches: 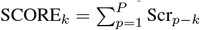. The most similar patient is selected by taking the one with the smallest distance.

### 3.3 Evaluation

#### 3.3.1 Mean average precision

In scenarios where retrieved entities (patches or patients) are assigned binary labels, we evaluate the retrieval performance using mean average precision (*mAP*), which is calculated as:

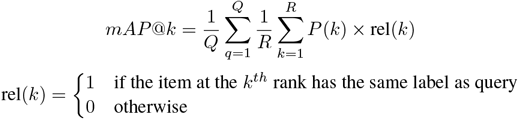

where *Q* is the total number of queries, *R* is the number of retrieved items for each query, *P* (*k*) is the precision at cut-off *k* in the list, and *rel*(*k*) is an indicator function equaling 1 if the item at rank *k* has a positive label, 0 otherwise.

#### 3.3.2 Euclidean distance

In the patch-level retrieval, both query and retrieved patches are assigned probability vectors representing the composition of different tissue structures. To evaluate its effectiveness, average Euclidean distances between the vectors are calculated and used as proxies for retrieval accuracy.

## 4 Results

In this section, we evaluate the performance of MIISS in various contexts, including patch and patient-level retrieval tasks in diabetic kidney disease (DKD), and patient-level retrieval between different cohorts of head and neck cancer (HNC) patients. We aimed to assess the robustness, effectiveness, and generalizability of the pipeline, as well as the performance of different featurizers and multichannel transformations. For more details on the datasets used, please refer to section A in the Appendix.

### 4.1 Case study - DKD kidney

We began with a qualitative case study in a DKD dataset generated with CODEX. We analyzed retrieval results obtained using a specified image patch in the DKD kidney study. As shown in Figure 2, the query patch identified by the red bounding box is extracted from a tissue section with severe DKD (class IIB). The patch is characterized by high concentrations of mucin 1 (MUC1) (magenta) and Collagen IV (yellow), suggesting distal tubule structures surrounded by highly fibrotic basement membranes. We utilized PLIP fusion model as the featurizer and identified the top 300 patches that are most similar to the specified query patch. Within the retrieved patches, we present examples from both the same region as the query patch and from different regions. Our observations reveal that these retrieved patches exhibit similar structures of distal tubules and fibrotic basement membranes, as suggested by the high concentrations of MUC1 and Collagen IV.

**Figure 2.**
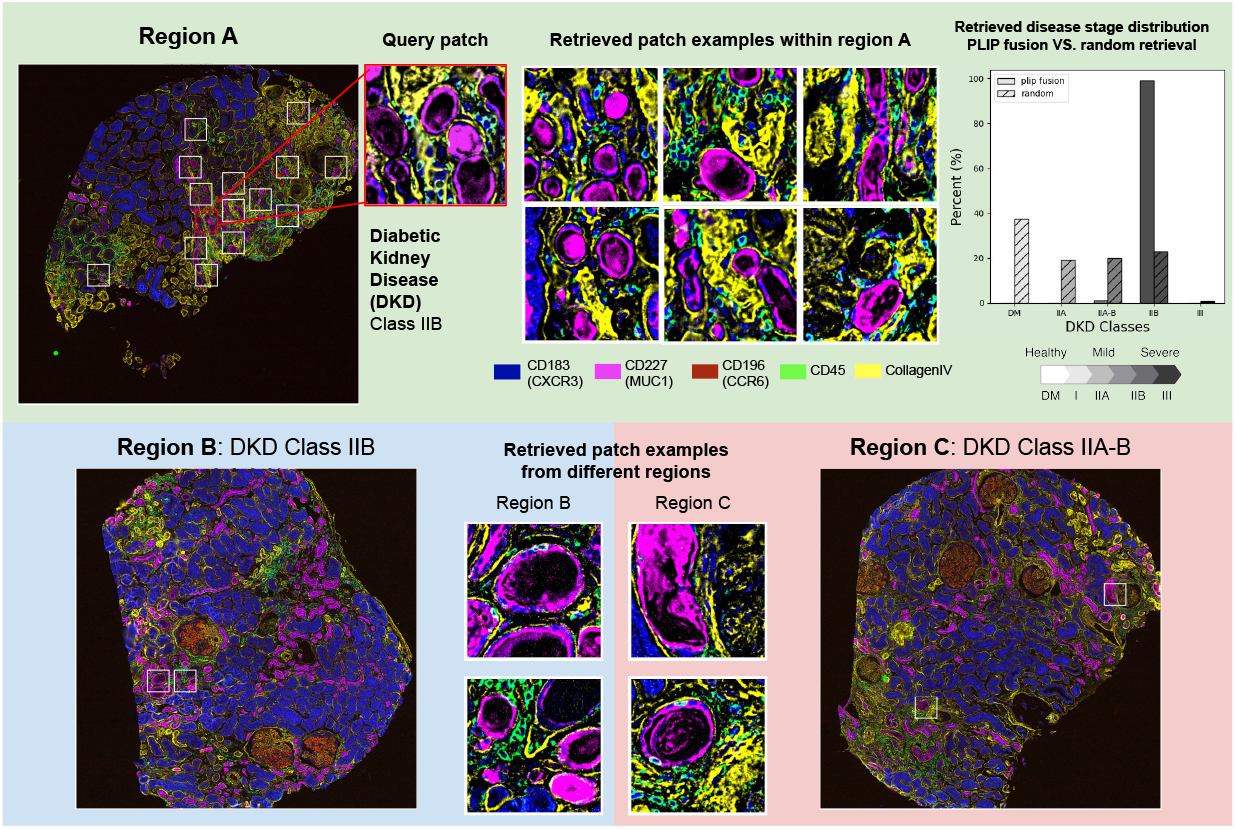
Case study on an example from the DKD Kidney study

To quantify the effectiveness of our retrieval method, we compared the distributions of DKD classes (of the patches) retrieved by PLIP fusion against random selection. The distribution of the randomly selected patches aligns well with the overall proportions of sample DKD classes, in which patches predominantly fall under the healthy kidney (DM) category. In contrast, the majority of the retrieved patches from PLIP fusion belong to the same DKD class as the query patch (IIB), with only a few exceptions from IIA-B intermediate samples that share similar pathological states as IIB samples. This comparison underscores the superior performance of our retrieval method. For a comprehensive evaluation, we also calculated the distributions of DKD classes retrieved by other featurizers and multi-channel transformations. Further details can be found in the Appendix.

### 4.2 Evaluate patch retrieval on kidney tissue structures

Patches from DKD kidney dataset contain diverse tissue structures. In the evaluation of patch-level retrieval, we assessed if the retrieved patches contained similar tissue structures as the query patches. To achieve this, we first obtained a probability vector for each patch representing the composition of its tissue structures, including proximal and distal tubules, glomeruli, blood vessels, and basement membrane. For each query patch, we subsequently computed the average *l*_2_ distance between its composition vector and the retrieved patches’ composition vectors. A lower distance indicates a higher similarity between the query and retrieved patches. As shown in Figure 3A, the feature fusion transformation consistently outperformed the random colorization alternative, with the notable exception of DinoV2. Among all the featurizers, PLIP fusion performed the best with the lowest average distance, suggesting that it excels in retrieving patches with similar tissue structure arrangements.

**Figure 3.**
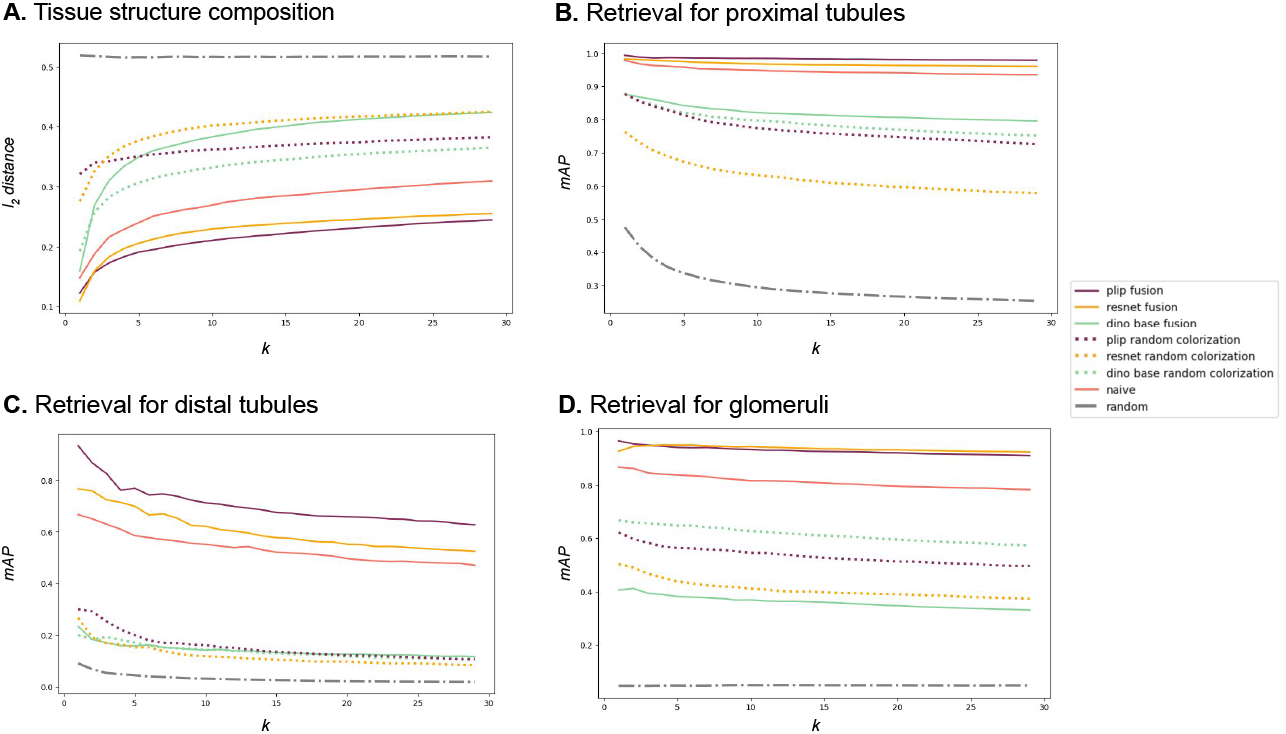
Patch level results based on cell type distribution

We further evaluated the performance of single-structure retrieval, in which patches that consist of over 90% of a single structure are used as queries. The mean average precision (*mAP*) metric is assessed based on the top-k retrieved patches, wherein a patch is deemed correctly retrieved if it contains more than 50% of the target structure; otherwise, it is classified as incorrectly retrieved.

The PLIP fusion model continued to excel, securing the highest mAP scores for the major tissue structures including proximal tubules and distal tubules (Figure 3B and C). For results on other tissues such as blood vessels and basement membrane, please see Figure 6 in the Appendix for more details.

In the retrieval task for glomeruli (Figure 3D), it came in a close second to the ResNet fusion model, with a negligible gap in performance. Both methods demonstrated robust performances in retrieving single-structure patches, achieving an mAP greater than 0.7 for the first 20 non-overlapping patches, which sets them apart from all other methods evaluated. Notably, the gap between PLIP fusion and the naive averaging method indicates that the spatial localization of biomarkers within patches provides additional structural information and assists with the correct retrieval of similar patches.

### 4.3 Evaluate patient retrieval on survival and HPV infection status

As demonstrated in the case study, the majority of the retrieved patches come from samples sharing the same disease state as the query patch. We aimed to further examine if the patient-level retrieval results are aligned in terms of pathological characteristics. We employed two head-and-neck cancer datasets to assess the MIISS framework. Both datasets are annotated with clinical data including the survival status and HPV infection status of patients. These annotations will be used to assess whether the retrieved patients are relevant to the query patient.

We first evaluated MIISS on the UPMC-HNC dataset by conducting a leave-one-patient-out cross validation. For each patient, we randomly selected 1,000 patches as queries and identified the top 20 most similar patients. The *mAP* metric is calculated on two clinical annotations of HPV status and survival status. As shown in Figure 4A, the majority of fusion methods, including the PLIP fusion model, exhibited strong performance and achieved high *mAP* scores across various cut-offs.

**Figure 4.**
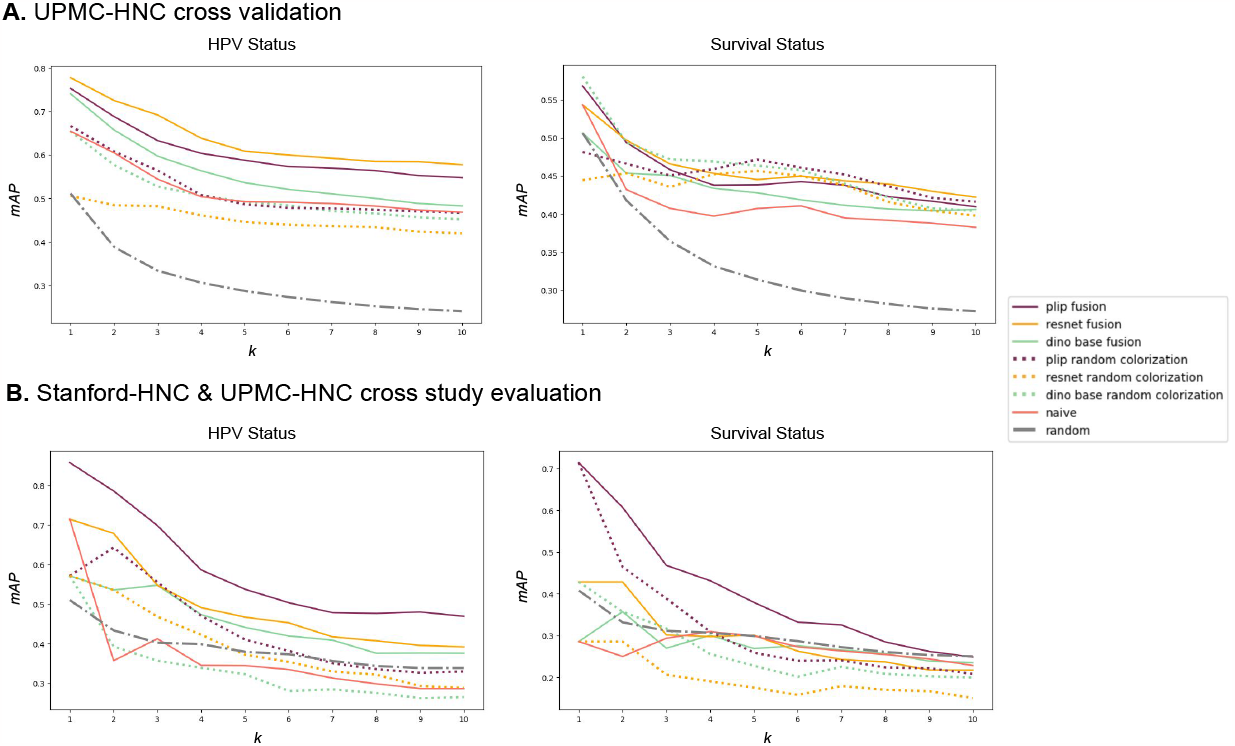
Patient outcome results on HPV status and survival status

**Figure 5.**
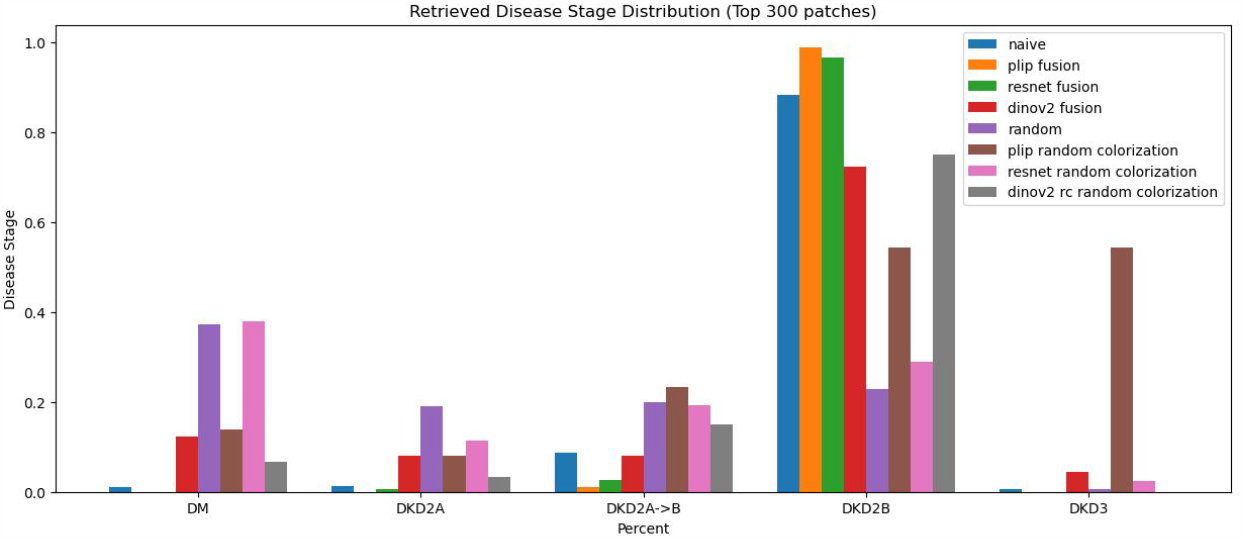
This figure compares the disease stage distribution between query patches and retrieved patches for all models.

**Figure 6.**
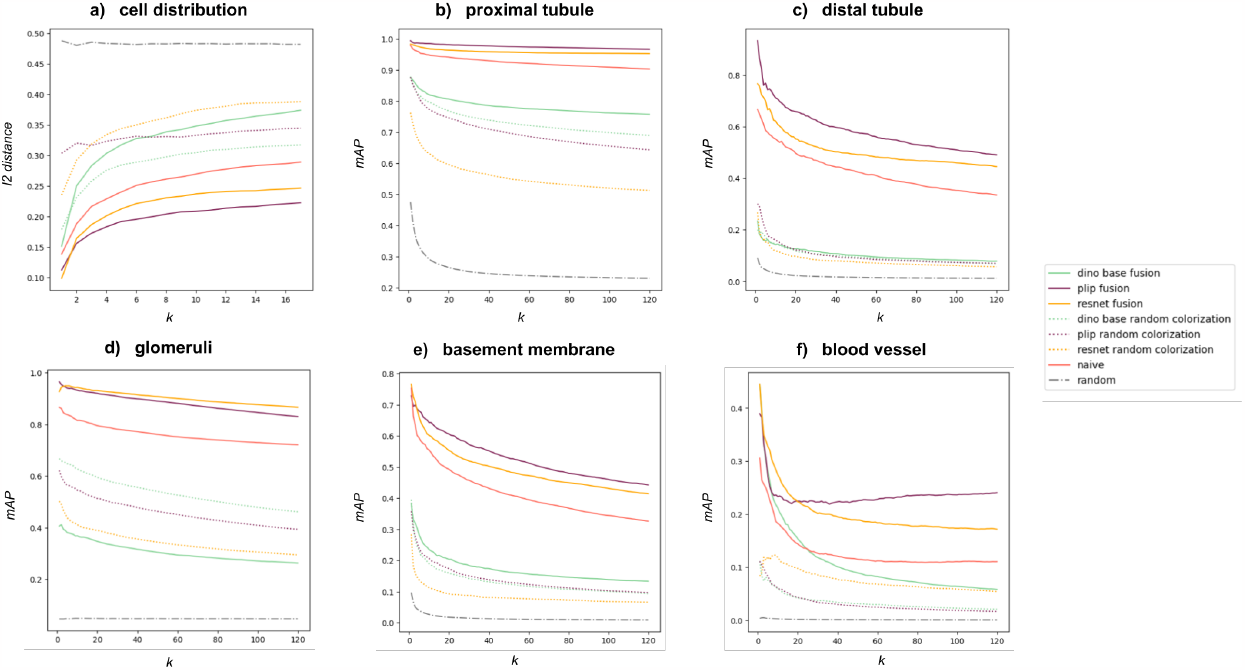
This figure compares the disease stage distribution between query patches and retrieved patches for all models.

To further simulate a realistic scenario, we performed a cross-study evaluation, where patients from the Stanford-HNC dataset were perceived as new entries with unknown HPV status and prognosis. By querying them against the UPMC-HNC dataset, we assessed if the MIISS retrieval framework can assist with diagnosis and prognosis predictions, thereby informing therapeutic decisions. Figure 4B presents the performances of all tested methods. PLIP fusion outperformed all alternatives by a notable margin, while the naive biomarker average method performed comparably with a random baseline. Such results highlight the importance of spatial localization of biomarker signals and morphological features in distinguishing tissues with different disease states or prognoses. Moreover, the robustness and adaptability of PLIP fusion are demonstrated by its superior performance across diverse datasets in both patch-level and patient-level retrieval tasks.

## 5 Discussion

Overall, we presented the MIISS framework, a novel image-to-image search pipeline for mIF imaging data. We leveraged advanced self-supervised learning and multimodal models for feature extraction and specifically evaluated the performance of DinoV2, ResNet and PLIP image encoders. We identified similar patches based on Euclidean distances between the query and retrieved patches, and employed an entropy-based scoring method to aggregate patch-level results at different granularities.

Our comprehensive evaluations on multiple datasets from different tissues demonstrated the robustness and effectiveness of the MIISS framework. At both patch and patient levels, PLIP fusion consistently outperformed other alternatives in retrieving similar patches or patients determined by tissue structure arrangements and disease states. Notably, PLIP fusion achieved an mAP@3 score of 0.7 for the cross-study retrieval task of HPV status, suggesting that a certain level of domain specificity substantially enhances the model’s generalizability. Moreover, it was not immediately evident that PLIP would generalize to the mIF case; however, we have proven its generalizability within the context of the mIF image search. This hints at the potential for a specialized pre-trained model tailored for mIF images.

While the MIISS framework shows promise, it is not without limitations. One drawback is the computational cost associated with feature generation, particularly when using the feature fusion technique. This could serve as a bottleneck for real-world applications. Therefore, more efficient alternatives, such as channel compression techniques, can be incorporated to optimize the feature generation process. Another limitation is that our current method doesn’t yet utilize all biomarker channels, as it selects intersected biomarkers across different studies. A future step could involve incorporating biomarkers that are less common. Despite these areas for improvement, the framework opens up new avenues for clinical benefits and research opportunities. Moving forward, we aim to broaden the applications of the MIISS framework to encompass a wider range of spatial omics modalities and disease types. We are also interested in developing a user-friendly interface to make the framework more accessible to pathologists and researchers, thereby facilitating its adoption in clinical and research settings.

## Appendix

### A Datasets

#### A.1 DKD Kidney

DKD kidney dataset contains 17 kidney cortical section samples from patients with diabetes and healthy kidneys (DM), diabetic kidney disease (DKD) classes IIA, IIB, IIA-B intermediate, and III. These samples were imaged and characterized using the CO-Detection by indexing (CODEX) platform on 21 protein biomarkers. We further obtained manual annotations of four major tissue structures: glomeruli, proximal tubules, distal tubules, and blood vessels. In this study, we analyzed if the retrieved patches contain the same tissue structures as the query patches.

#### A.2 UPMC-HNC

UPMC-HNC comprises 308 samples from 81 patients with head and neck squamous cell carcinomas. The tumor tissue samples were collected at the University of Pittsburgh Medical Center, and they were annotated with clinical data including survival status, disease recurrence, and HPV infection status of patients. In this study, we focused on how retrieval results reflect the survival status and HPV infection status of patients.

#### A.3 Stanford-HNC

Stanford-HNC is another head and neck squamous cell carcinoma cohort containing 26 samples from 7 patients. Samples were annotated with demographic information as well as cancer staging, HPV infection, survival and recurrence status. In this work, we used these samples to demonstrate the cross-study applicability of the similar patient search pipeline.

### B Retrieved disease stage distribution - All models

### C More patch level results

